# Capture, Confine, Characterize: High-Throughput Dielectrophoresis-Based Single-Cell Microfluidics Platform to Analyze Mammalian and Yeast Cells Using Raman Spectroscopy

**DOI:** 10.1101/2025.08.28.672773

**Authors:** Bum-Joon Jung, Allison Hohreiter, Dylan Cook, Linus Hansen, Sunny Taylor, Koseki J Kobayashi-Kirschvink, Joseph Chaiken, Supratik Guha, Anindita Basu

## Abstract

Single-cell analysis technologies are pivotal in unraveling complex biological mechanisms, yet existing platforms are often limited to sequencing-based end-point measurements, which fail to capture live cell dynamics. Here, we present a microfluidic– microelectronic device, the Microfluidic dielectrophoretic Arresting System (MiDAS) that employs dielectrophoresis (DEP) for high-throughput single-cell and droplet trapping in a compact array. We tested multiple trap geometries, including a 20 μm-diameter DEP trap for polymer microbeads, fungal and mammalian cells, and a 40 μm-diameter trap for water-in-oil droplets. The platform demonstrates broad sample compatibility, reliably immobilizing cells and beads of varying sizes. By integrating optical imaging and Raman spectroscopy, we enable rapid, non-destructive interrogation of individual cells with temporal resolution. We describe different modes of MiDAS operation to trap and manipulate single-cells or reverse emulsion droplets on demand, with applications in droplet microfluidics. Our MiDAS platform’s simple fabrication, robust performance, and broad compatibility with diverse sample types position it as a versatile tool with transformative potential for single-cell analysis, offering researchers an innovative approach to interrogate cellular dynamics with unprecedented precision and throughput.

## 1. Introduction

Single-cell analysis has enabled many advances in biology and medicine.^[1–9]^ However, single-cell techniques have been limited in the types of cells and samples that they can be applied to due to limitations in the available techniques for single-cell trapping and manipulation. Electric field assisted microwell separation^[10–15]^, droplet encapsulation16-18, hydrodynamic trapping^[19–24]^, and optical tweezers^[25, 26]^ are limited in their ability to accommodate cells of different sizes, high aspect ratio^[27]^, or diverse molecular compositions^[28]^, which prevent their broad application across different mammalian and microbial cells. Furthermore, cell recovery from methods such as antibody-coated microwell surface trapping^[29]^ can be difficult, limiting their ability to be combined with downstream analysis such as single-cell transcriptomics or proteomics. One technique that has promise to overcome these limitations is dielectrophoresis-based cell trapping.^[30–33]^

Dielectrophoresis (DEP) has been described previously as a method for trapping cells.^[34–36]^ It works by using patterned electrodes to apply an electric field gradient across a fluidic medium containing cells. Differential permittivity between the cell and the medium can lead to a net dielectrophoretic force that either: 1) directs cells to regions of low electric field (negative DEP force), or 2), directs them to regions of high electric field (positive DEP). Differences in controlling positive DEP (pDEP) and negative DEP (nDEP) processes have been described previously.^[37]^ Because the trapping mechanism relies on the electric field generated by patterned electrodes, cells can be easily trapped (and released) by simply turning on (and off) the voltage applied to the electrodes, thus flowing them off the MiDAS, as described in this study.

In this paper, we developed a simple DEP trap schema to trap and immobilize a range of entities, including cells, microbeads and reverse emulsion droplets. Our DEP trap design consists of two concentric metal electrode rings fabricated in a single layer, on glass or quartz substrates to generate an array of DEP traps in a single microfluidic channel. The design can be easily adjusted to change the trap size or the strength of the trapping force and scaled up to contain as many as ∼10,000 traps on a standard microscope slide. We simulated electric potential and field gradient of different DEP trap designs to optimize the design of electrode rings in silico and found good agreement between the simulation and MiDAS operation. We demonstrated different regimes of MiDAS operation to enable trapping of single or multiple cells by adjusting the size of the trap area (using device design), trap strength (using the amplitude and frequency of voltage applied to the electrodes), and flow parameters (flow velocity and cell concentration in the medium). Importantly, we leveraged the novel trap design and operation parameters to immobilize emulsion drops on DEP traps for the first time. We further demonstrate how the drops may be destabilized to release their contents, or merged with other drops, by adjusting the voltage and frequency of the DEP electrodes, indicating the MiDAS’s utility in droplet microfluidics applications. We show the MiDAS’s versatility by applying it to trap 5 μm and 10 μm polymer beads (as a proxy for cells), *Saccharomyces cerevisiae* (*S. cerevisiae*), *Candida albicans* (*C. albicans*), and mouse RAW 264.7 macrophage cells in device geometries compatible with bright-field and Raman microscopy, the latter of which has been used for non-destructive and time-resolved single-cell omics.38 Following on-chip imaging/spectroscopy, the trapped cells or droplets may be released from the traps and flowed off the MiDAS for downstream analysis, such as quantitative PCR, sequencing, or proteomics.

## 2. Results and Discussions

### 2.1. MiDAS Design and Working Principle

The MiDAS platform was designed to achieve high-throughput single-cell trapping through a scalable and straightforward fabrication process. **Figure 1a** illustrates an example MiDAS device design consisting of >10,000 single-cell traps. A photo of the device and magnified images of the DEP traps can be seen in **Figure 1b**. A single layer of Ti-Au electrodes is fabricated onto either a standard glass slide (to test and optimize DEP trap design) or a quartz wafer (for Raman spectroscopy measurements), using standard photolithography, E-beam, and lift-off methods.^[39, 40]^ The electrode layer consists of two concentric circular electrodes. The outer electrode ring is shaped like the letter “C” with a split opening to accommodate electrical connection to the inner electrode ring. The inner electrode ring is shaped like the letter “O” enclosing the DEP trap area. A PDMS microfluidic channel was created on top of the electrodes to allow sample, including cells, particles, or water-in-oil droplets, to flow over the electrodes at controlled flow rates. An AC voltage was applied to both the inner and outer electrodes to generate the electric field necessary for trapping. The transparent glass or quartz substrate provides light access to the cells, enabling optical and Raman microscopy from the bottom of the MiDAS for non-destructive and time-resolved characterization of cells or drops in DEP traps.

**Figure 1.**
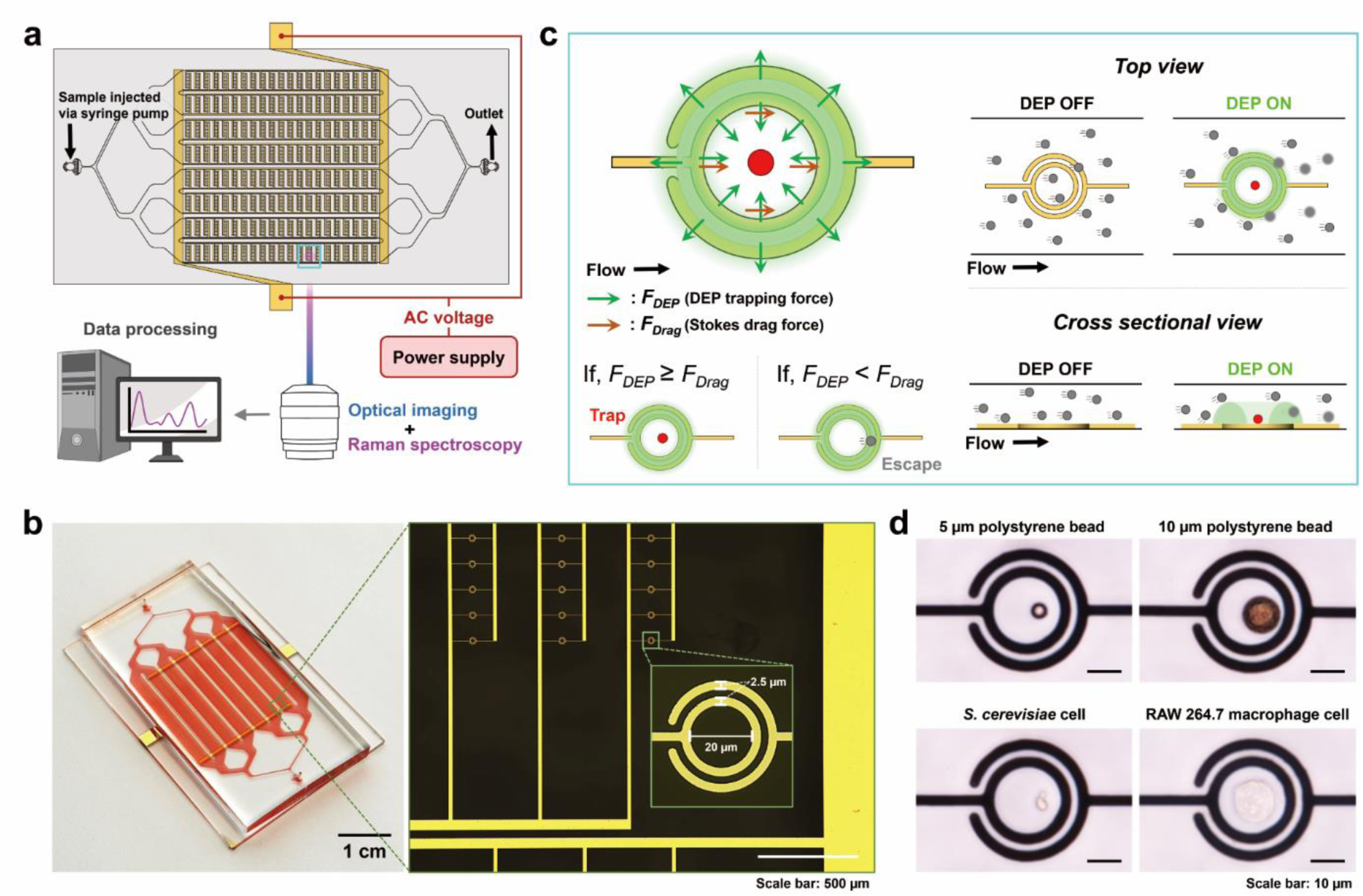
Microfluidic Dielectrophoretic Arresting System (MiDAS) platform for single-cell trapping and analysis. **(a)** Schematic illustration of the entire experimental setup, including Ti-Au electrodes fabricated on a glass/quartz substrate, PDMS channels for sample injection, and integrated bright-field, fluorescent, and Raman microscopy for real-time analysis. **(b)** Photograph of the assembled microfluidic-microelectronic device showing the PDMS microfluidic channel filled with red dye. Electrodes feature a 20 µm inner-ring diameter optimized for cell trapping and are composed of 150 nm Au electrode with a 10 nm Ti as an adhesion layer. **(c)** Schematic illustration of the negative dielectrophoresis (nDEP) working principle. Applied voltage generates a high electric field region (neon green band), creating an nDEP trap at the low-field minimum. Particles become trapped when the DEP force (green arrows) dominates over fluid drag force (orange arrows). Schematics compare particle flow with DEP field off ("DEP OFF") and activated ("DEP ON"). **(d)** Experimental demonstration of effective trapping for 5 µm and 10 µm polystyrene beads, yeast (*Saccharomyces cerevisiae*), and mammalian (RAW 264.7 macrophage) cells.

The working principle of the MiDAS (**Figure 1c**) relies on flow driven by a syringe pump to deliver microbeads (or cells), shown in red (or gray), to the vicinity of the trap and nDEP force to capture and immobilize the microbeads and cells inside the inner electrode ring. The time-averaged DEP force in the dipole approximation is^[41]^: 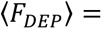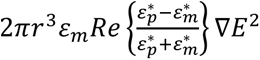, where 𝑟 is the particle radius and E is the electric field. The ε* with subscripts 𝑝 and 𝑚 are the complex dielectric permittivity of the particle/cell and medium, respectively. The nDEP force pulls the particle towards regions of low electric field strength. A high electric field gradient, illustrated as a green band, is formed between the inner and outer electrode rings when an AC voltage is applied. An electric field minimum is generated inside the inner electrode ring, creating the nDEP trap. In the absence of hydrodynamic flow, nDEP force is sufficient to trap cells/microbeads. However, in the presence of hydrodynamic flow, the nDEP force has to exceed the hydrodynamic force, 𝐹_𝐷𝑟𝑎𝑔_ = −6𝜋𝑟𝜂𝑣 (given by Stokes’ law), where 𝑟 is the particle radius, 𝜂 is the fluid viscosity, and 𝑣 the particle velocity, to be able to hold the particle in the nDEP trap. Trapped particles, represented in red, are held on the surface of the substrate due to dominant DEP forces (green arrows), while untrapped particles, shown in gray, are carried away by hydrodynamic force (orange arrows) in the direction of the fluid flow.

When the DEP field was deactivated ("DEP OFF"), the particles move/diffuse freely inside the DEP trapping region. In contrast, with the field activated ("DEP ON") at low applied voltage, a single particle or cell can be immobilized in the trap. By increasing the applied voltage, more than one cell can be captured per trap. Thus, by modulating the applied voltage, it is possible to reach the desired cell loading per trap.

The assembled MiDAS (**Figure 1b, left**) shows a design consisting of >10,000 DEP traps. A red dye was introduced into the PDMS channels to enhance visibility of the microfluidic channel atop the electrode layer fabricated in gold (**Figure 1b, right**). To ease the burden of fabrication (by fitting multiple devices on the same mask) and test the working principles of the traps under multiple conditions, we used MiDAS containing arrays of 5x5 traps per device. However, the MiDAS can easily accommodate >10,000 traps on a standard microscope slide to facilitate high-throughput operations. CAD files for DEP trap designs with 25 and 10,000 traps per device are provided as Supplementary Material.

We fabricated and tested two different DEP traps with inner electrode rings of 20 μm and 40 μm diameters, henceforth referred to as 20 μm traps and 40 μm traps, respectively, using the same fabrication process for both. The electrode lines are 2.5 μm in width and 160 nm in height, with a spacing of 2.5 μm between the electrode lines. Each electrode ring consists of a 150 nm thick layer of Au, deposited atop a 10 nm thick Ti layer added first to improve Au adhesion.^[42]^ **Figure 1b**, right (inset) shows a 20 μm trap with Au electrodes, where the DEP trap region inside the 20 μm diameter inner electrode ring is slightly larger than a typical mammalian cell^[43]^, to ensure effective trapping. The spacing between the inner and outer electrode rings is 2.5 μm, equal to the width of the gold electrodes. We optimized the operation parameters (cell/bead concentration, flow rate, applied voltage and frequency) to trap and immobilize single polystyrene microbeads (5 μm and 10 μm diameter), *S. cerevisiae* S288C (yeast), and RAW 264.7 macrophage (mammalian) cells in the 20 μm traps (**Figure 1d**). The 40-μm trap design was altered proportionally, such that the width of the spacing between the gold electrodes, as well as the width of each electrode in the 40 μm trap is 5 μm.

### 2.2. Characterizing MiDAS Operation *via* Simulation and Experiments on Cells and Microbeads

To validate the electric field configuration, numerical simulations of the electric potential (V) and gradient of the electric field squared (∇𝐸^2^ ∝ 𝐹_𝐷𝐸𝑃_) were performed on the 20 μm trap design for different applied voltage (5, 10 and 20 V peak-to-peak (Vpp)) and frequency (1 - 20 MHz, in 2 MHz increments) (**Figure 2a-c**). Our simulations show that a voltage differential between the inner and outer ring (**Figure 2a, left**) generates a region of high ∇𝐸^2^ between the rings, while the region within the inner ring exhibits lower ∇𝐸^2^, where the nDEP trap region is formed (**Figure 2a, right**). Due to the asymmetry in the DEP trap design (stemming from the split outer electrode ring), there is a small asymmetry in the electric potential and ∇𝐸^2^. This asymmetry is easily seen in the cross-sectional view of the electric potential in the xz (but not yz) plane (**Figure 2b**), with inset figures illustrating the voltage in 3D. Resulting from this asymmetry, the region of lowest ∇𝐸^2^ shifts slightly off-center in the DEP trap, away from the split opening of the outer electrode ring. Arrows indicating ∇𝐸^2^ (**Figure 2c**) at different applied voltages (5, 10, and 20 Vpp) and 1 MHz frequency confirm vertical confinement, demonstrating that the DEP trap can effectively confine particles in x, y and z directions. The simulation results agree well with experimental observation (**Figure S1a**), verifying the MiDAS’s reliability in confining particles under the applied AC voltage. Consistent with the dielectrophoretic field simulations (**Figure 2c, S1a**), the particles are immobilized slightly off-center in the DEP trap. Specifically, the particles are immobilized away from the center of the DEP trap, repelled by the high ∇𝐸^2^ at the split outer electrode. A split outer electrode design with sharp edges (**Figure S1b**) can create an additional nDEP trap right by the split opening. Smoothing the edges of the split electrode can bypass the formation of this additional (albeit weaker) trap. For simplicity, our simulations focus on mapping the overall DEP force field only, and do not incorporate permittivity of the particles, or drag force on the particles under flow, which we approximate to be uniform across the MiDAS, under volume-driven flow at low flow rate.

**Figure 2.**
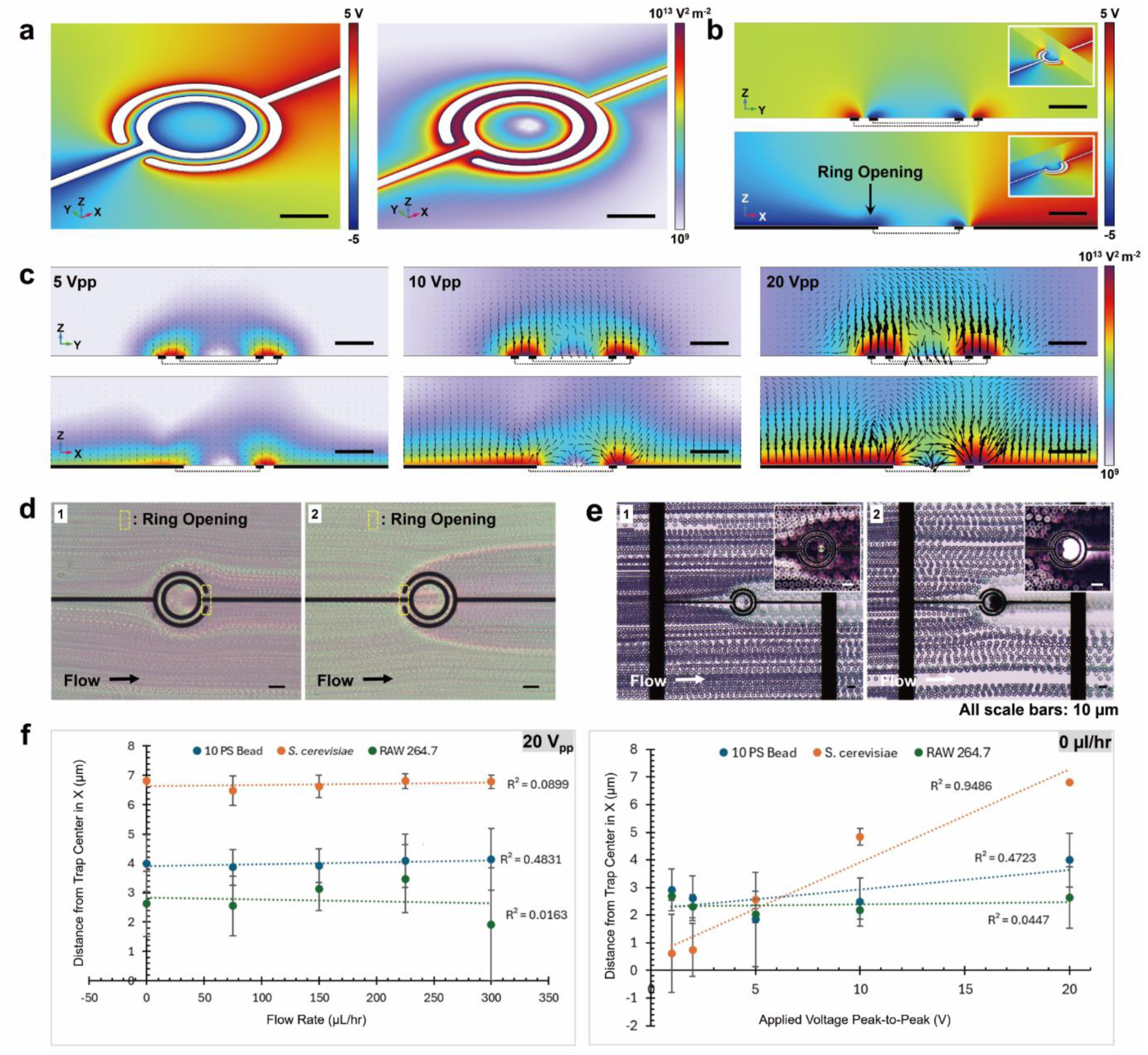
Electric field COMSOL simulations and particle trapping analysis under various flow conditions. **(a-c)** COMSOL simulations illustrating the electric field distribution in a 20 µm DEP trap. **(a)** 3D angled view showing electric potential (left) and electric field norm squared (right) showing high electric field (∼10^13^ V^2^/m^2^) between electrode rings and low field (∼10^9^ V^2^/m^2^) at the inner trap where nDEP region is formed. **(b)** Cross-sectional views highlighting electric potential which shows asymmetry of the electric potential on a Z-X plane due to the ring opening. Inset figures represent the respective cross-sectional planes in 3D views. **(c)** Detailed cross-sectional electric field norm squared (surface) and the negative gradient of the squared magnitude of the electric field (arrows) profiles at varying voltages (5, 10, and 20 Vpp), confirming vertical confinement. The arrows indicate the direction of the nDEP force in logarithmic vector form. **(d)** Flow trajectories of *S. cerevisiae* cells influenced by ring opening orientation relative to flow: (1) opening opposite flow and (2) opening facing flow shown by maximum intensity plots from 500 image stacks on a 5% BSA coated 40 µm DEP traps. **(e)** 5 µm polystyrene beads trapping behaviors under balanced conditions (E1, single-bead trapping) and DEP force-dominant conditions (E2, multiple-beads trapping) shown by minimum intensity plots from 300 image stacks; insets: enlarged standard deviation plots from the same stacks. **(f)** Quantitative analysis of particle displacement relative to trap center as a function of flow rate at fixed applied voltage, 20 Vpp (left) and applied voltage at fixed flow rate, no flow (right) for 10 µm polystyrene beads, RAW 264.7 cells, and *S. cerevisiae* cells.

The velocity profiles of particles (cells or beads) flowing past the traps was different, depending on the orientation of the ring opening relative to direction of flow; **Figure 2d** shows the (maximum) intensity projection of image stacks generated from video of *S. cerevisiae* cells flowed in and against the ring opening. With the ring opening facing away from oncoming flow (**Figure 2d1**), the particles were accelerated along their trajectories, flowing in a semi-circular path that keeps roughly with the dimensions of the trap before returning to their previous trajectory. When the ring opening is facing the oncoming flow (**Figure 2d2**), oncoming cells are either trapped or pushed away from the trap. This behavior can be visualized in the electric field simulations (**Figure 2a, right**). Cells are trapped in the low electric field region inside the inner electrode ring (light purple colored central region), consistent with nDEP trapping. Conversely, cells approaching the regions of significantly higher electric field gradients around the electrode rings (regions marked by red, indicating elevated field intensity, ∼10¹³V²/m²) experience repulsive forces, resulting in their being deflected or pushed away from the trap. Regardless of flow direction relative to the ring opening, trapped cells accumulate inside the inner ring, away from the outer ring opening. We used the flow-DEP trap conformation in **Figure 2d2** for this study.

Aggregate particle trajectories under different trapping regimes are illustrated in **Figure 2e1, 2e2**, using minimum intensity projections of images in a video (**Movie S1 and S2**). **Figure 2e1** instantiates the regime where F_DEP_ ∼ F_Drag_. At higher voltages, where F_DEP_ > F_Drag_, multiple particles can be trapped simultaneously in a single trap (**Figure 2e2**). The insets show particles in DEP traps using intensity projection (standard deviation) of videos used in **Figure 2e1, 2e2**.

Other factors that influence the precise location of the trapped particle are velocity, size, and dielectric property of the particles (**Figure 2f**), which were not captured in our simulations. These parameters were empirically tested. To locate the precise position of entrapped particles in the DEP trap under different flow velocity, particle types or suspension media, we conducted imaging experiments at various flow rates and voltages on 10 µm polystyrene beads, RAW 264.7 macrophages and *S. cerevisiae* cells (suspended in 1X PBS). We measured the center of the beads/cells and the center of the DEP trap (using NIH ImageJ) to calculate the displacement of the bead/cell center from the center of the trap in the direction of flow (**Figure 2f**), repeated across three different microbeads (and cells) per condition. We observed noticeable displacement of the particles from the trap center with increasing applied voltage and flow rate. We posit that the displacement does not continue to increase with increasing flow rates because at low flow rates the DEP force on the particle within the trap is relatively small. Thus, the particle can be pushed around in the trap by the applied flow until it reaches a region of the trap with sufficient force to hold the particle in place. At higher flow rates, the particle is pushed away from the center of the trap, where it encounters rapidly increasing DEP force and therefore unable to be displaced further. The DEP traps are unable to hold particles at flow rates > 350 µL/hr, as F_Drag_ exceeds F_DEP_.

*S. cerevisiae* cells, which are smaller than the 10 µm microbeads and mammalian cells, show the largest displacement from the trap center as function of flow rate and applied voltage (**Figure 2f**). The small size of the *S. cerevisiae* cells also meant that variations in cell placement from the trap center due to differences in flow rate and applied voltage could be observed more easily, and these variations have been plotted as kymographs (**Figure S2a,b**). These kymographs confirm that the trapped particles are not stuck irreversibly to the glass/quartz slide upon trapping but instead exhibit slight rocking and rotational motions over time.

We developed two methods for cell trapping with our DEP trap design. In the first method, under continuous flow, we turned on the applied voltage to activate the DEP field when a single particle (cell or microbead) passed directly above the inner electrode ring of the trap. At low flow rate (*e.g.*, 75 µL/hr), a weaker DEP field was sufficient to trap and immobilize particles using relatively low applied voltage (*e.g.* 5 Vpp). However, this method lacked scalability, requiring each trap to be monitored and activated transiently to capture single particles.

Our second method of trapping can overcome the need to monitor individual traps. First, we filled the microfluidic channel (*e.g.* 75 µL/hr) with particles (microbeads or cells) at high concentration (*e.g.* 600,000 particles/mL), while the DEP field was off. After the microfluidic channel was filled, the flow was stopped and the particles were allowed to come to rest. Under these conditions, there was negligible fluid drag, causing the particles to sediment onto the glass/ quartz surface. We then applied a small voltage (∼ 1 Vpp) to activate the DEP field. This induced a small DEP force, drawing nearby cells or particles into DEP trap. Once the DEP field was activated, we gently increased the flow rate to wash away particles outside the nDEP region, leaving only one trapped particle. This method was most effective when the nDEP trap size was slightly smaller than the particle size. If particles were much smaller than the nDEP trap size, they often clustered together and behaved collectively as a single object. Conversely, if the nDEP trap region was smaller than an individual particle, only a single particle remained tightly trapped in the nDEP trap region, while particles that were weakly trapped by the DEP force were washed off.

To evaluate the potential impact of nDEP trapping on cell viability, *S. cerevisiae* cells suspended in 1X PBS were subjected to electric field (10 Vpp, 1 MHz) and stained with 0.1% Methylene Blue (IBI Scientific. No. IB74050) at 1:1 volume ratio. Cells in PBS and trapped in the DEP field were stained within ∼ 41 seconds but took longer than 3 mins to stain when 0.5% BSA was added to the suspension. This assessment provides insight into cell membrane integrity under electric field. Further studies are warranted to optimize experimental parameters for individual cells to enhance their viability/ membrane porosity for live-cell analyses.

### 2.3. MiDAS Application to Trap or Merge Droplets

We also tested the MiDAS for utility with droplet microfluidics, as many current single-cell techniques rely upon water-in-oil droplets not only to isolate cells^[3]^, but to confine a cell’s secretome or lysate.^[44]^ We employed the 40 µm DEP trap with a 50 µm height microfluidic channel to immobilize or merge reverse emulsion droplets, depending on applied voltage, frequency and flow rate. Drops of different diameters: >75 µm (orange, **Figure 3a**; yellow, **Figure S3b**) and <30 µm (blue, **Figure 3a, S3b**), containing 1X PBS and suspended in an inert carrier oil, were tested. The permittivity of 1X PBS at pH 7.4 (inner phase) is similar to that of water^[45]^, which is ∼80 F/m.^[46]^ In contrast, the permittivity of the droplet generation oil (CAS. No. 297730-93-9) comprising the continuous outer phase is ∼6 F/m.^[47]^ Thus, we observed positive DEP force on the droplets, drawing and immobilizing them to the thin annular spacing between the inner and outer electrode rings. We found 75 µm droplets entrapped across the 40-µm DEP trap (**Figure 3a, left**). In contrast, the smaller droplets were trapped in the annular space between the electrode rings (**Figure 3b, left**).

**Figure 3.**
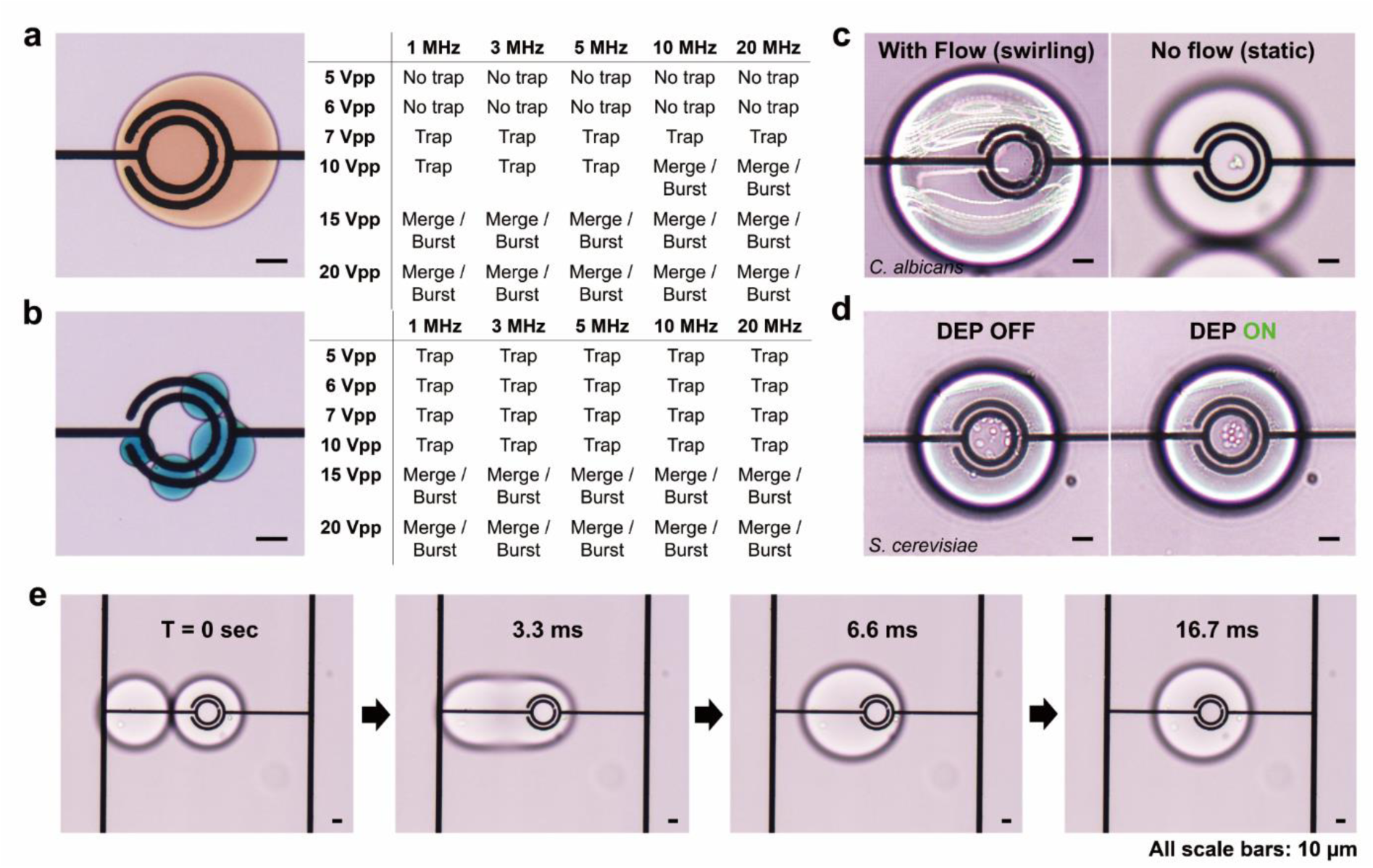
Application of the DEP device to water-in-oil emulsion droplets trapping and merging. **(a-b)** Images showing trapping performance for **(a)** 75 µm and **(b)** 30 µm and smaller droplets, along with corresponding summary tables **(a-b, right)** detailing results under varying voltage and frequency conditions. These illustrate size-dependent trapping efficiency and dependency on microfluidic channel height. **(c-d)** Cells in droplets trapping: **(c)** *C. albicans* cells and **(d)** *S. cerevisiae* cells within droplets, demonstrating cell and droplet immobilization at the DEP trap upon DEP activation **(c-d, right)**, compared to free cell movement under flow conditions **(c, left)** or no flow and DEP OFF conditions **(d, left)**. **(e)** Time-lapse images showing *C. albicans* cells in droplet merging with at 20 Vpp, 1 MHz, under no-flow conditions.

Since the height of microfluidic channel (50 µm) was larger than the smaller drops (< 30 µm), and aqueous droplets being less dense than the carrier oil^[47]^, the smaller drops flowed above the region of the DEP force field, often escaping the trap. By restricting our microfluidic channel height to 25 µm, we demonstrated that the droplets < 30 µm were also effectively captured (**Figure 3b**), with multiple droplets being captured along the annular space with high ∇𝐸^2^. Various voltage and frequency trapping conditions were tested, and their trapping efficiencies are reported for both larger droplets (**Figure 3a, right**) and smaller droplets (**Figure 3b, right**). The dependency of droplet trapping effect on channel height is worth noting; droplet-based single-cell applications can often benefit from size selection of droplets to aid experimental reproducibility. We also noted chaining of droplets, a common effect of droplets in applied electric field.^[48, 49]^

While the droplet trapping operates in the pDEP regime, larger droplets, which cover the entire trap area, can function as single DEP traps for particles suspended within. To demonstrate this principle, we trapped ∼50 to 75 µm droplets containing yeast cells (suspended in PBS) encapsulated in oil on MiDAS with 20 µm DEP traps (height = 50 µm), where the cells were drawn to the center of the DEP trap (**Figure 3c, 3d**). When the DEP field was OFF, the droplets stayed pinned to the DEP traps but spinning in the direction of flow at flow rates below 500 µL/hr. The droplet spinning, in turn, caused the encapsulated *C. albicans* cell to spin within the droplets; **Figure 3c**, left shows the maximum intensity projection of images from a video of this phenomenon (**Movie S3 and S4**). However, increasing the flow rate above this threshold resulted in droplets getting unstuck from the traps, thereby allowing retrieval of encapsulated cells for downstream applications, such as RNA-seq. Conversely, with the flow stopped and DEP turned ON, the *C. albicans* cell (within the drop) was confined to the center of DEP trap (**Figure 3c, right**). Similarly, multiple *S. cerevisiae* cells would stay dispersed within a droplet, when the DEP field Lastly, but importantly, we are able to merge droplets immobilized on a DEP trap by increasing the applied voltage on the trap (**Figure 3e**). The voltage and frequency dependence of droplet trapping/ merging for 75 & 30 µm droplets are summarized in **Figure 3a, 3b** right, respectively. The timescale of two droplets merging is shown in **Figure 3e**. This important functionality can be useful to deliver reagents (e.g., drug, lysis agent) to cells encapsulated in drops and immobilized on DEP traps, for on-chip optical/spectroscopic characterization. At higher voltages (≥15 Vpp), we observed droplet merging and shrinking/electrolysis effects (**Figure 3a, 3b right**). We assume that polar surfactants within the droplet generation oil, which typically function to stabilize the droplets during formation, facilitated their degradation at high voltages, causing the droplets to shrink and eventually burst.

### 2.4. MiDAS Application for Optical Microscopy and Raman Spectroscopy

Finally, we demonstrated the MiDAS’s utility for optical microscopy and Raman spectroscopy applications (**Figure 4**). Previous studies have combined Raman microscopy with single-cell transcriptomics to monitor changes in live cells in real-time without the need for chemical labels^[38]^. The configuration of our MiDAS-Raman setup is shown (**Figure 4a**). Here, we image and collect Raman spectra using a Renishaw Virsa Raman analyzer through the bottom of the microfluidic device consisting of 525 µm thick quartz^[38]^. We obtained Raman spectra (3 accumulations at 532 nm, 62 mW and 1 s) from single mammalian (mouse RAW macrophage) cells in MiDAS with the DEP turned off.^[50, 51]^ The process of Raman spectra collection using the MiDAS device is demonstrated in **Figure 4b**. The raw Raman spectra were corrected for sample auto-fluorescence using a baseline-correction method^[52]^ of 101-7 moving-average for each raw spectrum. The baseline corrected spectra for 13 different RAW cells collected on-device with the DEP OFF is shown in **Figure 4c, left**, and the cell-to-cell Pearson correlations are shown in **Figure 4c, right**. While we were able to capture single-cell heterogeneity between individual macrophage cells using Pearson correlation of their baseline-corrected spectra, we noted significant background signal from the PDMS elastomer that was used to construct the device microchannel.

**Figure 4.**
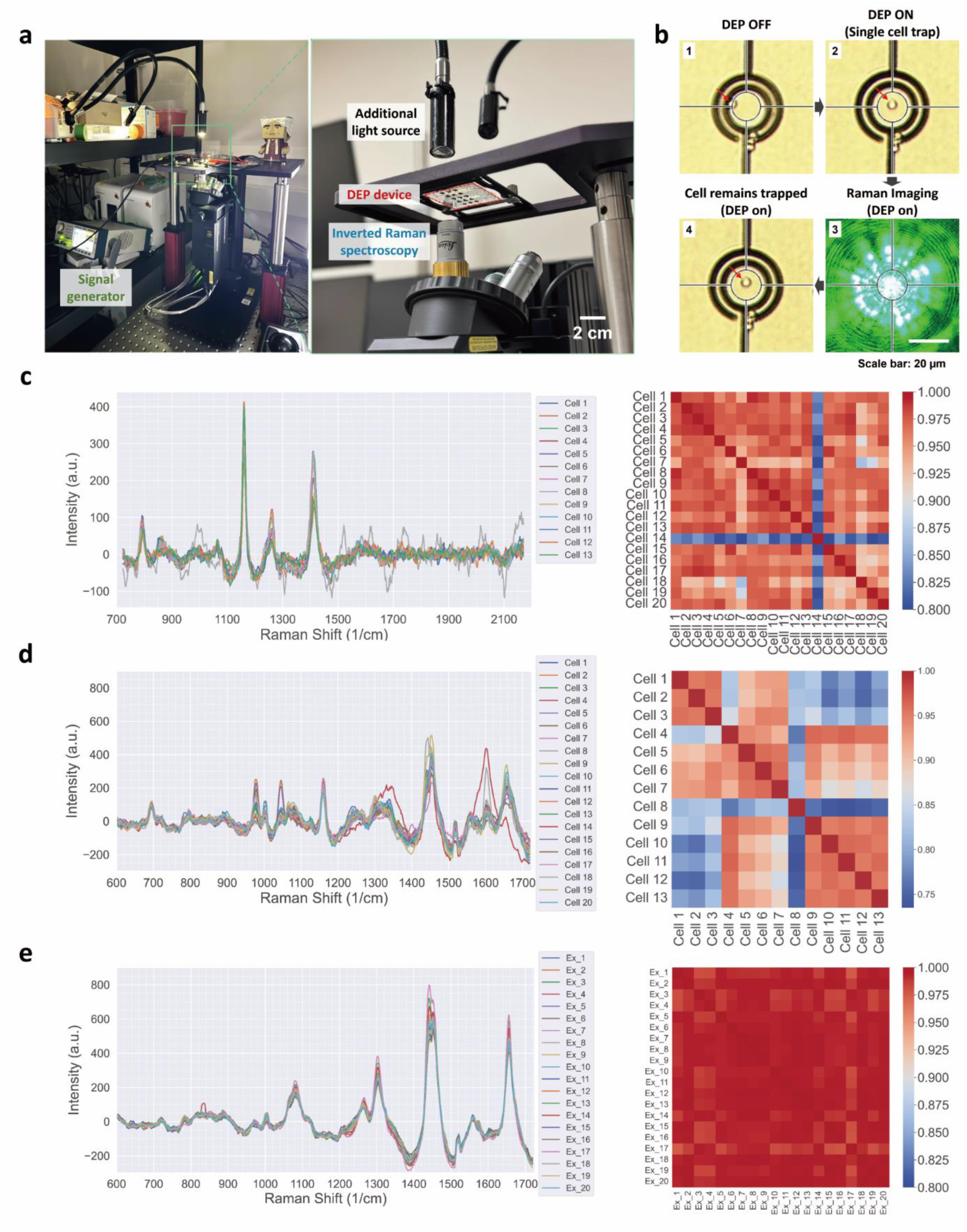
On-chip single-cell analysis using the integrated MiDAS-Raman spectroscopy platform. **(a)** Photograph of the experimental setup, showing the MiDAS device mounted on an inverted microscope coupled with a Raman analyzer and a signal generator. **(b)** Single-cell capture and analysis of a *C. albicans* cell: (1) A cell approaching an nDEP trap (DEP OFF), (2) DEP activation capturing a single-cell, (3) a focused laser used for Raman spectroscopy of a trapped cell, and (4) the cell after Raman spectroscopy. **(c)** Raman spectral analysis of individual RAW 264.7 macrophage cells on the MiDAS: The plot shows baseline-corrected spectra from 13 different cells (left), and the Pearson correlation matrix (right) between cells. **(d)** Raman spectra from 20 different *S. cerevisiae* cells dried on a substrate (left), with the corresponding correlation matrix (right). **(e)** Twenty Raman spectra acquired from a single *S. cerevisiae* cell repeatedly (left) and their corresponding Pearson correlation matrix (right).

To mitigate the contribution of PDMS elastomer on the cell spectra, we constructed a microfluidic device as before, but now consisting of a quartz chamber sitting atop the DEP traps that were fabricated on a 525 µm thick quartz substrate (as before). Using this quartz-on-quartz device, we were able to capture single-cell spectra of RAW macrophages with well-defined Raman spectral peaks (**Figure S4a, left**), obtained after baseline-correction. The Pearson correlation of Raman spectral shift between ∼700-2200 cm-1 for 21 different RAW macrophage cells sampled at random using this quartz-on-quartz configuration was ∼0.85 (**Figure S4a, right**). Next, we examined the correlation of repeated Raman spectral measurements on three single macrophage cells (**Figure S4b, left**). These measurements showed a slightly higher Pearson correlation for measurements of the same cell (∼0.88, **Figure S4b, right**). However, with the DEP turned on, the Pearson correlation between Raman spectra obtained under identical settings was lower, e.g., between different macrophage cells (∼0.78) or the same cell, measured repeatedly (∼0.82) (**Figure S4c**). This may be due to cell electroporation caused by the applied electric field.^[52, 53]^ By keeping the Raman spectra acquisition time short (∼1 s/cell), it is possible to minimize the electroporation effects on our spectra. As comparison, the Pearson correlation of Raman spectra obtained from the background PBS suspension media (1X PBS) in this configuration was ∼0.95, which being of uniform composition, is higher (**Figure S4b, right**).

Raman spectroscopy of microbial cells at single-cell resolution is challenging because of their low signal.^[54, 55]^ To investigate if Raman spectroscopy has the ability to resolve single-cell level heterogeneity in microbial cells, we performed benchmarking experiments on dried *S. cerevisiae* cells using a Horiba LabRAM Raman confocal microscope (532 nm, ∼3 mW, and 2 accumulations at 30 s per acquisition). Given the lower laser power, the spectra were acquired over longer duration (30 s) and baseline-corrected as before, using the 101-7 moving average scheme^[56]^ to factor our sample auto-fluorescence. We compared the spectra of 20 different *S. cerevisiae* cells dried on a quartz coverslip (**Figure 4d, left**) with the spectra of a single-cell taken 20 times (**Figure 4e, left**) and their Pearson correlations (**Figure 4d, e, right**, respectively). The average Pearson correlation between multiple Raman spectra from the same cell was higher (∼0.99) than the average Pearson correlation between the spectra from different *S. cerevisiae* cells (∼0.95).

Challenges of spontaneous Raman spectroscopy of biological samples include weak signal^[57–60]^ and sample auto-fluorescence. Surface Enhanced Raman Spectroscopy (SERS) can boost Raman spectral signal, which can be especially beneficial to microbial cells. To leverage this phenomenon, we added a 150 nm thin layer of gold onto a quartz substrate and dried *S. cerevisiae* cells onto the gold-coated surface to enhance their Raman spectroscopy signal. For comparison, a control experiment was conducted using cells dried directly onto bare quartz. As shown in **Figure S5a**, the spectra from cells on the gold layer exhibit significantly enhanced intensity (∼10x) compared to cell spectra on bare quartz. After baseline correction, we note that the Raman spectral peaks from cells dried on gold-coated quartz aligned with the peaks from cells dried on bare quartz substrate (**Figure S5b**). To leverage this phenomenon, a thin layer of gold may be added inside the inner electrode ring (but electrically isolated from it), to amplify signal from cells/samples with weak Raman spectra.

## 3. Conclusion

The MiDAS’s ability to handle diverse sample types while maintaining the ability to capture, immobilize, monitor and release single particles on demand underscores its robustness and versatility. We demonstrated the ability to trap different particle types of different size, including polymer microbeads, yeasts and mammalian cells. Additionally, the MiDAS’s ability to capture cells encapsulated in emulsion droplets or merge droplets containing cells on-command while maintaining nDEP capture of cells represents a new utility for DEP systems with implications for experiments requiring reagent delivery to cells, e.g., mixing cells isolated in aqueous droplets and immobilized on DEP traps (for spectroscopy) with lysis cocktails, chemokines, or fresh media. In addition, the MiDAS architecture is compatible with in-line Raman spectroscopy and optical microscopy with utility in single-cell profiling. Finally, because the number, size or strength of the DEP traps on a MiDAS is easily modified and scalable, it can fill a new niche for high-throughput single-cell analysis on-chip and droplet merging on-demand.

### Limitations and outlook

The current design of MiDAS platform for single-cell analysis using Raman spectroscopy has several limitations that present opportunities for future improvement: A key challenge is the significant background signal from the PDMS elastomer used to construct the microfluidic channel, which interfere with cellular Raman spectra. Future design of the device can address this by replacing PDMS with a material with lower auto-fluorescence or Raman emission spectra, to construct microfluidic channels.

Another challenge is low signal-to-noise ratio (SNR) of Raman spectra of microbial cells. To improve SNR, employing a thinner glass or quartz substrate, or using an alternative material with minimal overlap in Raman spectra at relevant excitation wavelengths would be beneficial. Another strategy to enhance SNR, particularly for samples with weak Raman signal like microbial cells, is to employ surface enhanced Raman scattering (SERS) phenomena by adding an Au pad within the inner DEP ring, where the cell is trapped, to amplify its Raman spectra.^[61, 62]^ Finally, we observed that the electric field used for DEP trapping can likely induce electroporation of cells. Therefore, a critical future task will be to identify experimental conditions that minimize electroporation, thereby enhancing the accuracy and reliability of time-resolved analyses of single-cells.

## 4. Experimental Section/Methods

### MiDAS Device Fabrication

The DEP electrodes were fabricated utilizing conventional photolithography, soft lithography, E-beam metal deposition, and lift-off techniques. Before photoresist deposition on a glass slide (Fisher Scientific; 3″ × 2″ × 1.2 mm thick) or 4-inch quartz wafer (University Wafers, #2298), HMDS vapor treatment (YES-58TA Vacuum Bake) was applied to improve adhesion between the substrate and upcoming photoresist. A layer of AZ nLOF 2020 photoresist (Microchem) was spun at 3000 rpm for 30 s, resulting in about a 2.0 μm film thickness, and soft baked at 110 °C on a hot plate for 60 s. Then, the laser pattern was applied to pattern the electrode design under Heidelberg direct write lithography machine (Heidelberg MLA150) using i-line sensitive (365 nm) with the exposure energy of 250 mJ/cm2. Subsequently, the photoresist was developed using the AZ 300 MIF developer (Microchem) for 30 sec with gentle agitation. Before metal layer coating, rinse the chip with Deionized water (DI water) for a minute and blow dry with nitrogen spray guns. Afterwards, the chip was treated with oxygen plasma for the removal of resist residues and contaminants from the surface. 10 nm thick titanium and 150 nm thick gold were successively deposited by electron beam evaporator (Angstrom EvoVac). Finally, the slides were soaked in an 80 °C NMP solution overnight to lift off the sacrificial photoresist and the overlying metal layer. PDMS microfluidic channels^[63]^ were fabricated by spin-coating SU-8 3025 (Kayaku) on 4-inch silicon wafers (University Wafers, #452) at 1200 rpm for 30 s with acceleration of 300 rpm/second, resulting in a microchannel layer ∼50 μm thick device used to entrap microfluidic droplets. Spin-coating SU-8 at 3000 rpm for 30 s with acceleration of 300 rpm/second resulted in a microchannel layer ∼25 μm thick used to entrap cells or microbeads. The SU-8 was soft baked on a 95 °C hot plate for 15 min for both of the microchannel types. The design for the channel was patterned on the SU-8 using a Heidelberg direct write lithography machine (Heidelberg MLA150) and subsequently hard baked at 65°C for 1 min and 95 °C for 5 min. This process was identical for both microchannel types. The pattern was then developed using SU-8 developer for 5∼8 min with a gentle agitation. To cast the microchannel layer from the SU-8 mold^[63]^, PDMS (Krayden Dow Sylgard 184 Silicone Elastometer Kit) was mixed in a 10:1 elastomer:cross-linker ratio in a THINKY AR-100 centrifugal mixer for mixing (30 s) and degassing (30 s), and then poured directly onto the SU-8 mold. The PDMS was then further degassed gently in a vacuum chamber and then placed in a 65 °C oven overnight to set. To assemble the device, the set PDMS channels were cut and peeled from the SU-8 mold using a scalpel and pressed gently into place on the gold electrodes patterned on the glass (or quartz) slide, leaving the electrode connector pads uncovered (**Figure 1c**). The PDMS layer was aligned by eye with the DEP traps fabricated on glass or quartz slides and held in place by simply pressing against the slide. Two conductive pins were soldered to each of the two electrode connector pads.

Prior to MiDAS operation, the PDMS channels were treated to prepare the devices for either direct application of cells in 1X PBS or water-in-oil droplets. For direct application of cells in 1X PBS to the MiDAS, the channel was treated using 5% BSA in 1X PBS overnight at 4 °C to prevent sticking of the cells to the glass (or quartz) substrate or PDMS.

### Cell Culture and Preparation for MiDAS Usage

*S. cerevisiae* strain S288C and *C. albicans* strain SC5314 were grown in Yeast Peptone Dextrose (YPD) (MP Biomedicals) on a Thermo Scientific MaxQ 4000 shaker set at 30 °C with rotation at 300 rpm. Cells were collected from YPD liquid cultures, pelleted by centrifugation at 700 xg for 5 minutes, washed 3 times with 1X PBS, and diluted to an appropriate working concentration for imaging prior to droplet encapsulation or direct application on the MiDAS. Murine RAW 264.7 macrophages were cultured in complete medium (Dulbecco’s Modified Eagle Medium, high glucose (Gibco) supplemented with 10% FBS (Sigma-Aldrich) and 1% Penicillin/Streptomycin (Gibco) in an incubator kept at 37 °C with 5% CO_2_. Cells were cultured in T25 flasks and used below 20 passages. RAW 264.7 cells were allowed to grow to ∼60% confluency, and then collected by removing the complete medium and gently washing with 1X PBS followed by a 5-minute treatment with 10% trypsin-EDTA at 37 °C, which was then quenched with complete medium followed by cell scraping to retrieve the cells from the flask. The cells were then collected by brief centrifugation at 300 xg for 5 minutes. If being prepared for droplet encapsulation or direct application to the MiDAS, after centrifugation the RAW 264.7 cells were resuspended in 1X PBS and then collected by centrifugation at 300 xg for 5 minutes. They were then washed an additional 2 times with 1X PBS for a total of 3 washes with 1X PBS and diluted to an appropriate working concentration for imaging prior to droplet encapsulation or direct application on the MiDAS.

### Bead Preparation for MiDAS Usage

Two different sized polystyrene microparticles, 4.95 μm (Bangs Laboratories. Inc.) and 10 μm (Molecular Probes™ CML Latex Beads) diameter were used to observe microbead trapping in MiDAS. Microparticles from the stock solutions were resuspended in 1X PBS and then collected by centrifugation at 300 xg for 5 minutes. They were then washed an additional 2 times with 1X PBS for a total of 3 washes with 1X PBS and diluted to an appropriate particle concentration in the range of 0.01−0.05 w/v%. In addition, to prevent particle aggregation, a 1% of non-ionic surfactant (Tween 20, Sigma-Aldrich) was added to the particle suspension.

### Droplet Generation

The aqueous phase (cells prepared as described above, or beads in 1X PBS) was loaded into a 3 mL syringe. EvaGreen droplet generation oil (Bio Rad) was used without further purification and loaded into a 10 mL syringe. Both syringes were capped using 25G needles attached to PE2 microfluidic tubing (Scientific Commodities) and injected into the microfluidic channel using two KDS910 syringe pumps (KD Scientific). Water-in-oil droplets of various sizes were then generated using a 30 µm droplet co-flow device fabricated using standard soft lithography techniques.^[63]^ The droplet generation channel was briefly treated with Aquapel (PGW Auto Glass, LLC.) for 30 s and dried using filtered air. This increases the hydrophobicity of the glass side of the channel and aids droplet stability. The aqueous phase was flowed through the cell inlet, the oil phase flowed through the oil inlet, and the bead inlet was blocked off as it was not in use. The parameters used for droplet generation are as follows: for 75 μm droplets, a 75 μm wide channel with ∼ 70 μm depth was used with flow rates of 4 mL/hr for oil and 2 mL/hr for the aqueous phase. For 30 um droplets, a 30 μm wide channel was used with flow rates of 2 mL/hr for oil and 0.5 mL/hr for the aqueous phase. Droplets were collected in a 50 mL falcon tube and immediately reloaded into a 3 mL syringe to be injected into the MiDAS.

### MiDAS Operation and Optimization

For direct usage on suspended cells or beads: The electronic components of the MiDAS were controlled by a signal generator (BK Precision 4064 B) which was connected to the conductive pins on the device via alligator clips. Cells or beads in 1X PBS were loaded into a 3 mL syringe and capped using 25G needles attached to PE2 microfluidic tubing as mentioned previously. The cells/beads in 1X PBS were then flowed into the microfluidic channel of the MiDAS using the same pumps and pump software as described previously. Once media containing cells/beads had perfused the microfluidic channel, we collected videos of the cell/bead behavior within the traps under various MiDAS operation parameters to determine optimal operating conditions. Namely, the applied peak-to-peak voltage on the traps was varied across the range of our signal generator, at 1, 2, 5, 10, 15 and 20 V. The flow rate was also swept from 0 to 300 μL/hr in increments of 75 μL/hr. The frequency of the MiDAS was held at 1 MHz. We noticed steady erosion of the electrodes at frequency ≤ 1 MHz. For usage with water-in-oil droplets: We tested the MiDAS in both a standard orientation (electrode on the bottom, and PDMS channel on the top) and an inverted orientation (electrode on the top, and PDMS channel on the bottom). As illustrated in **Figure S3**, the standard orientation led to droplet merging without bursting (**Figure S3a**), whereas the inverted orientation caused droplets to burst and wet the glass surface (**Figure S3b** red arrows). In the standard orientation, PBS droplets float in oil while contacting only the hydrophobic PDMS surfaces; in contrast, in the inverted orientation, the floating PBS droplets come into contact with the hydrophilic glass surface. We believe this wetting occurs because, upon droplet merging, the droplet contacts the hydrophilic glass, resulting in droplet bursting and wetting the glass. For all (cell, bead or droplet) operating modes, brightfield images were acquired using an inverted brightfield microscope (Olympus CKX53) and a Teldyne Chameleon 3 FLIR camera.

### Modeling of the Electric Field using COMSOL

As the dielectrophoretic force on cells and particles to trap them is proportional to the gradient of the electric field squared, we simulated the following parameters across the microfluidic channel, using COMSOL Multiphysics v6.0: electric potential (V), displayed on a cut plane using a rainbow color scheme that scales linearly; norm of the electric field squared, ‘es.normE^2’ (V^2^/m^2^) and shown in prism color scheme that scales logarithmically; gradient of the squared electric field, ∇𝐸^2^ ‘∇(es.normE^2)’ (V^2^/m^3^), indicated by arrows, the size of which scale logarithmically. AutoCAD (Autodesk, Inc.) files containing the electrode designs were converted into closed loops, using the ‘Region’ function and exported as separate .dxf files such that there were no nested loops in any file. The .dxf files were imported into COMSOL under ‘Geometry’ into a work plane and two separate electrodes consisting of the inner and outer rings were reconstructed in COMSOL using ‘Boolean’ operation. The work plane containing the electrodes was extruded to desired thickness (*e.g.*, 0.2 μm) and assigned the material properties of gold (Au) from the COMSOL ‘Materials’ library. Above the electrodes, an additional work plane with a thickness of 25 μm was created to simulate the microfluidic channel layer, within which a ’Water, liquid’ layer (simulating PBS with a relative permittivity of 80 F/m) was defined. The geometry, combining the three work planes, was meshed using Physics-controlled mesh with element size set to ‘Extremely fine’. The ‘Electrostatics’ Physics were used to simulate the electric potential and the gradient of the electric field squared, being proportional to F_DEP_. The gradient of the squared electric field, ∇𝐸^2^ (arrows) indicates the direction of nDEP in the xz plane, which has the expression of x-component: -d(es.normE^2^,x) and y-component: -d(es.normE^2^,z) and unit, kg^2^*m/(S^6^*A^2^) = V^2^/m^3^. Frequency-dependent voltages of 5, 10, and 20 V were applied in the range from 1 - 20 MHz.

### Image Analysis

Images were analyzed using Fiji.^[64]^ Maximum intensity projections of the particles’ position in the traps were generated by using the Z-project tool on video stacks that consisted of ∼400 frames. From these max intensity projections, we manually drew circles using the oval tool over the object and the trap center. Then, we used the Measurement function to measure the centroid of both the captured object and the trap center. We found the X displacement relative to the center of the trap by subtracting the trap’s center X position from the object’s center X position. We did this for each condition using a total of 3 particles, from which an average displacement in X and standard deviation was calculated. Kymographs were generated to analyze X-direction object movement over time within the traps under each condition using the raw video stacks previously mentioned. Briefly, a mask of the trap was created by the Thresholding tool. From here, the inner ring trap object was selected and its centroid found. The Y-position of the centroid was used to draw a horizontal line across the midpoint of the trap, which was used to run the kymograph function (pixel width = 51) on the image stack.

### Raman Spectroscopy

Initial measurements were performed on dried yeast *(S. cerevisiae*) cells, using a confocal Raman microscope (HORIBA LabRAM HR Evolution) at UChicago MRSEC core facility. Yeast cells were washed in deionized water and dried on a quartz coverslip (TED PELLA, INC. No. 26016) at low concentrations such that individual cells could be interrogated by Raman spectroscopy. The cells were excited at 532 nm at ∼3 mW and Raman spectra was acquired for 30 s, and 2 acquisitions, using a 600 lines/mm grating. Raman spectra of RAW macrophages in 1X PBS on a MiDAS (through quartz) were acquired using a Renishaw Virsa Raman Analyzer at 532 nm and ∼62 mW excitation.

Briefly, the cells were washed three times in 1X PBS to limit signal from added molecules in culture media. Raman spectra were acquired with 3 accumulations of 10 s each, using a 50x ELWD objective and a 2400 lines/mm grating. Measurements were performed through either the PDMS-based channel lid or the 525 µm quartz substrate.

### Raman Spectroscopy data analysis

Raw emission spectra of each cell were exported as text files. Raman emission spectra between 501-1725 cm-1 (*S. cerevisiae*) and 723-2170 cm-1 (RAW macrophage) were compared, using Pearson correlation. Baseline-correction was performed^[56]^ to minimize the contribution of auto-fluorescence signal on the Raman spectral shifts; Pearson correlation of the baseline-corrected spectra was visualized as heatmaps for easy comparison, using Python 3.8.

## Supporting information

CAD Files

## Supporting Information

Supporting Information is available from the Wiley Online Library or from the author.

## Acknowledgements

This work made use of the Pritzker Nanofabrication Facility, part of the Pritzker School of Molecular Engineering at the University of Chicago, which receives support from Soft and Hybrid Nanotechnology Experimental (SHyNE) Resource (NSF ECCS-2025633), a node of the National Science Foundation’s National Nanotechnology Coordinated Infrastructure [RRID: SCR_022955]. This work was supported by the Duchossois Family Institute Fellowship (BJ.J.) and NIH DP2AI158157 grant (A.B.).

## Conflict of Interest

The authors declare no conflict of interest.

## Data Availability Statement

The data that support the findings of this study are available from the corresponding author upon reasonable request.

## Funding

Soft and Hybrid Nanotechnology Experimental (SHyNE) Resource (NSF ECCS-2025633), a node of the National Science Foundation’s National Nanotechnology Coordinated Infrastructure [RRID: SCR_022955]. This work was supported by the Duchossois Family Institute Fellowship and NIH DP2AI158157

## Notes

### Competing Interest Statement

The authors have declared no competing interest.

