## Supplementary Information for "Capture, Confine, Characterize: High-Throughput Dielectrophoresis-Based Single-Cell Microfluidics Platform to Analyze Mammalian and Yeast Cells Using Raman Spectroscopy"

Bum-Joon Jung, Supratik Guha
Pritzker School of Molecular Engineering, University of Chicago, Chicago, IL 60637, USA

Dylan Cook
Committee on Genetics, Genomics, and Systems Biology, University of Chicago, Chicago, IL 60637, USA

Linus Hansen, Sunny Taylor
The College, University of Chicago, Chicago, IL 60637, USA

Joseph Chaiken
Department of Chemistry, Syracuse University, Syracuse, NY 13244, USA

**Supporting Information**

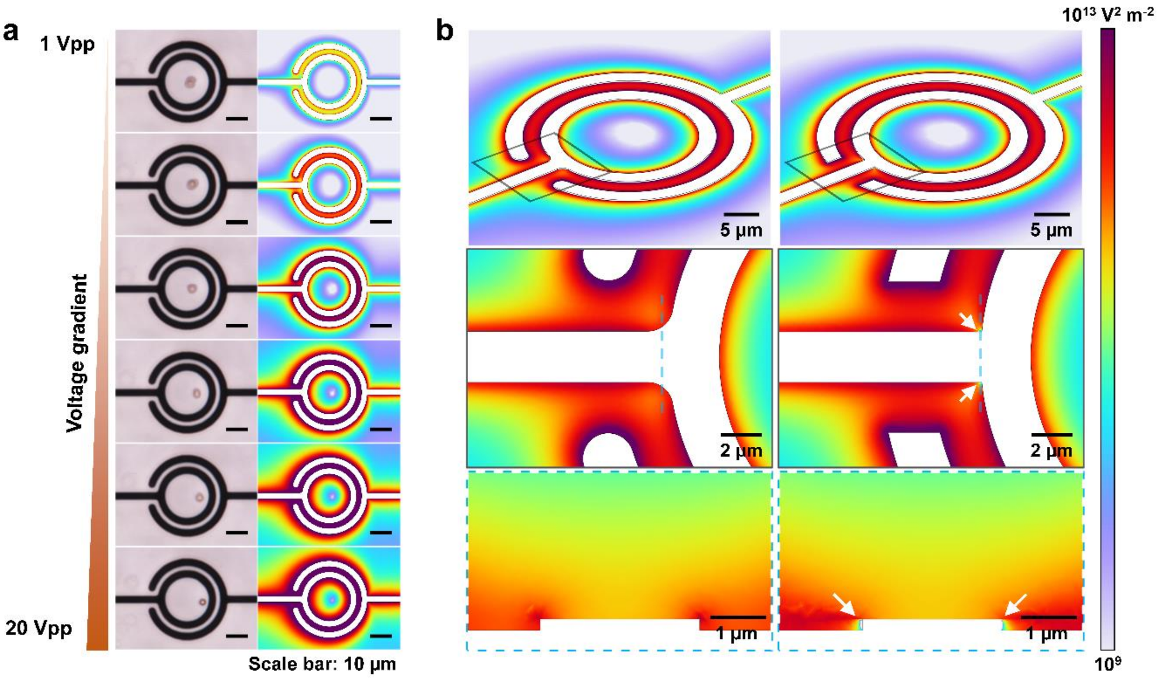


**Figure S1.** Experimental and COMSOL simulations of electric field distribution and particle trapping behavior. **(a)** Comparison of experimental images (left column) and corresponding COMSOL simulations of the electric field norm squared (right column) at increasing voltages (1 to 20 Vpp), demonstrating particle trapping slightly off-center due to asymmetric electrode design. **(b)** Detailed simulations highlighting the impact of electrode edge geometry; sharp edges (right column) create unintended smaller nDEP trapping regions (white arrows), potentially trapping particles outside the intended inner ring area. Smooth edge (left column) electrode designs eliminate these unintended trapping sites.

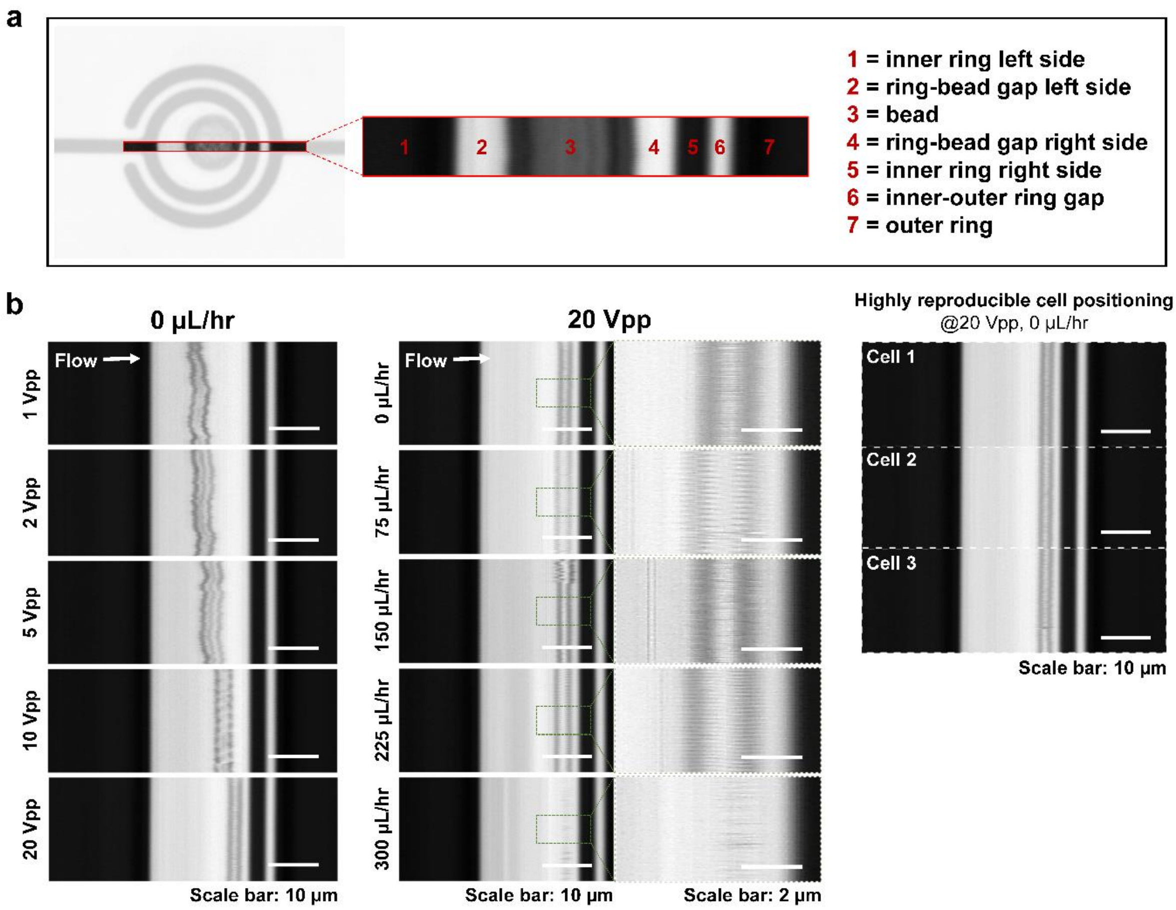


**Figure S2.** Kymographs of objects trapped in 20 µm DEP trap. **(a)** Kymograph interpretation key demonstrated on a 10 µm bead. The inset shows a representative 1209x51 pixel snapshot, which is taken from each image in a video to generate the kymographs. Within this snapshot, the dark left-most band (1) is the left side of the inner ring. The light left-most band (2) is the gap between the bead and the left side of the ring. The middle gray band (3) is the bead/trapped object. The first light band on the right of the bead (4) is the gap between the bead and the inner ring right edge. The next dark band (5) is the right side of the inner ring. The next right light band (6) is the gap between the inner and outer rings, and the final dark band (7) is the outer ring. These snapshots are stacked on top of one another according to their order in the video to generate the final kymograph. **(b)** Kymographs of a trapped *S. cerevisiae* cell under various conditions: at different applied voltages with no flow (left); at different flow rates with a fixed voltage (20 Vpp), where the periodic patterns (highlighted) indicate cell rotation (middle); and for three different cells (replicates) under identical static conditions (20 Vpp, 0 µL/hr) to show trapping reproducibility (right).

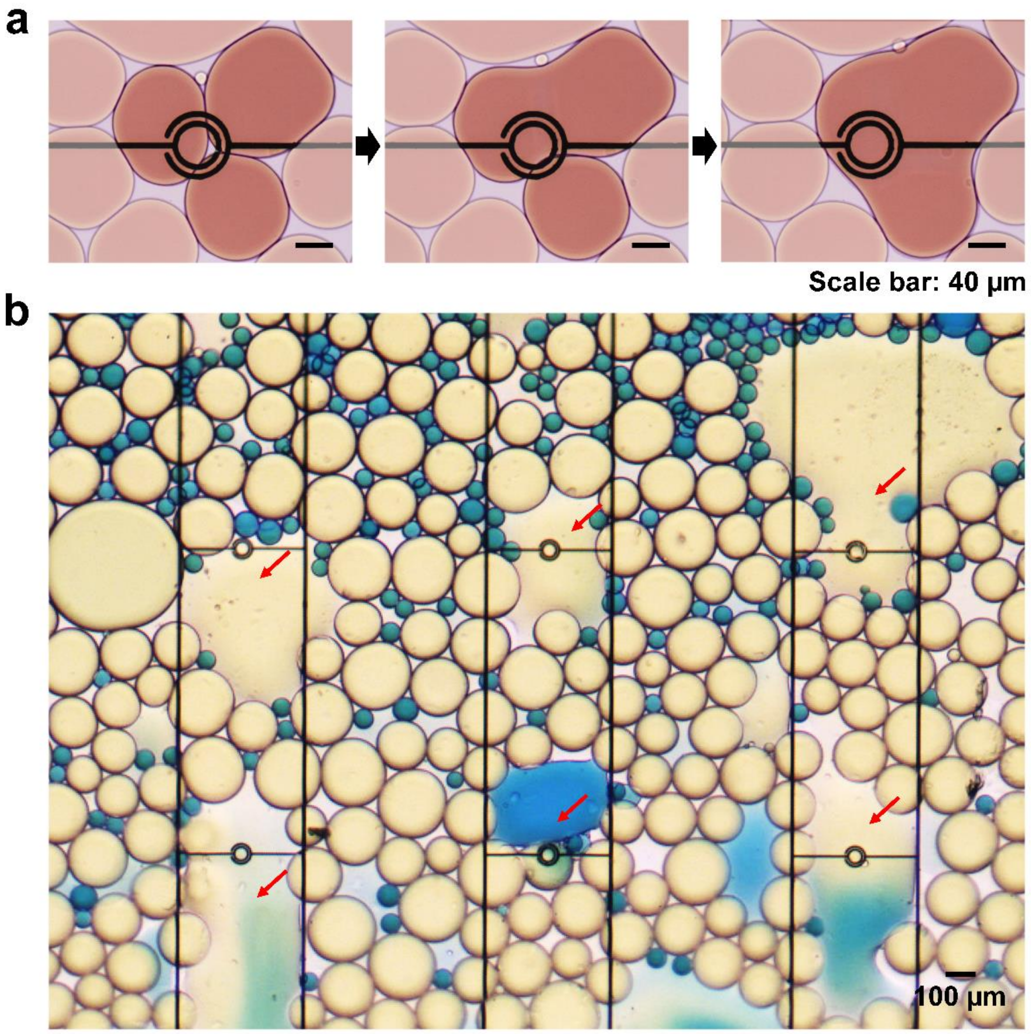


**Figure S3.** Distinguishing between droplet merging and bursting based on device orientation. **(a)** Controlled droplet merging is achieved in a standard microchannel configuration where the electrodes are on the bottom glass substrate. In this setup, droplets primarily contact the hydrophobic PDMS walls, leading to stable merging without bursting. **(b)** In contrast, when the microchannel is inverted (electrodes on top), droplets burst and wet the hydrophilic glass surface, as indicated by the red arrows. This is because the less dense aqueous droplets rise through the oil and contact the now-overhead hydrophilic glass, causing them to rupture upon merging.

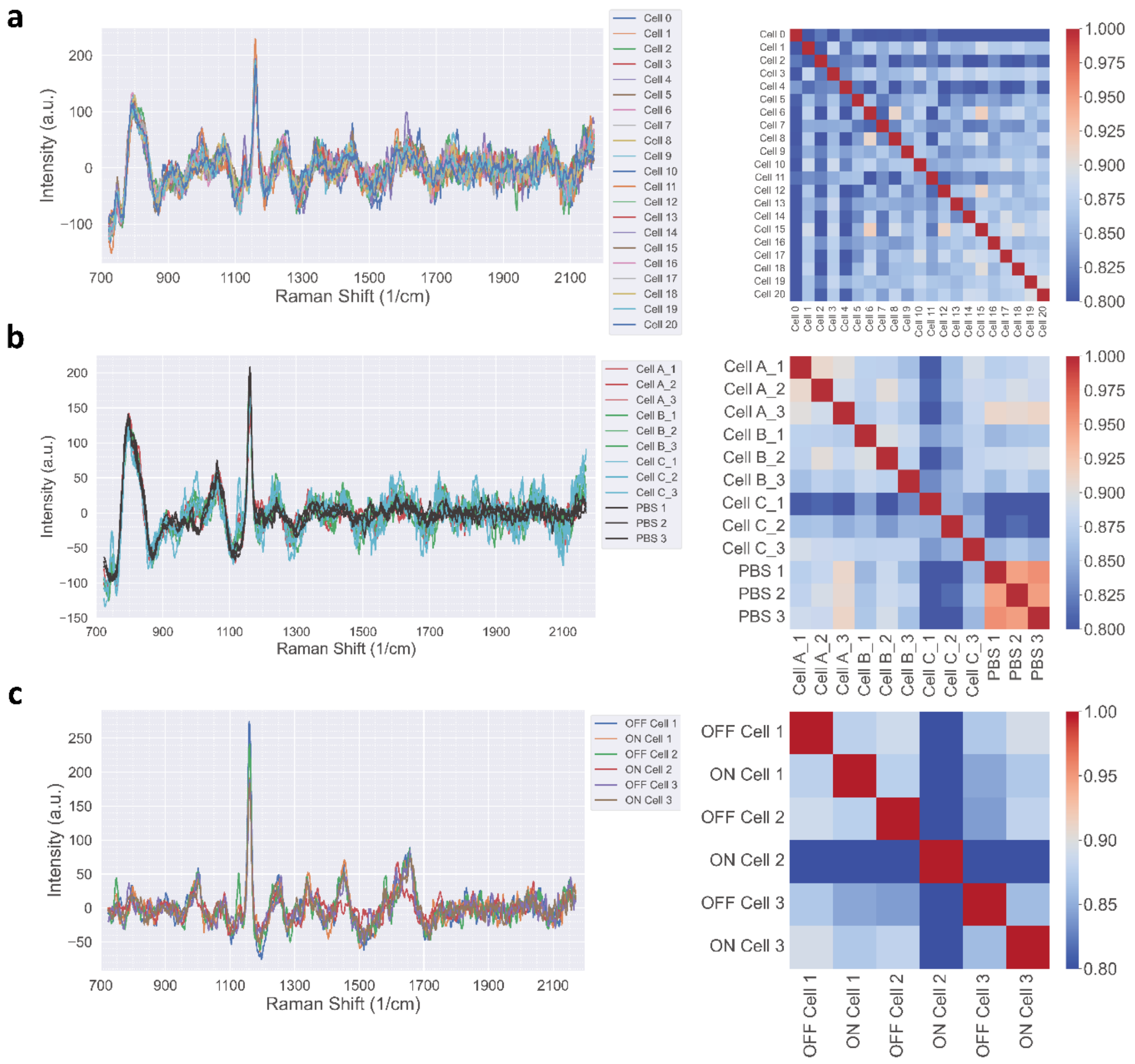


**Figure S4.** Raman spectral analysis of single macrophage cells using a quartz-on-quartz microfluidic device to minimize background signal. **(a)** Baseline-corrected Raman spectra from 21 individual RAW macrophage cells (left) and corresponding Pearson correlation matrix (right). **(b)** Repeated spectral measurements and the Pearson correlation matrix from three different single macrophage cells **(a, b, c)** and 1X PBS background (3 repeats each; right). (b) Raman spectra from three individual cells with the DEP field turned OFF and ON (left) and the correlation matrix of their spectra.

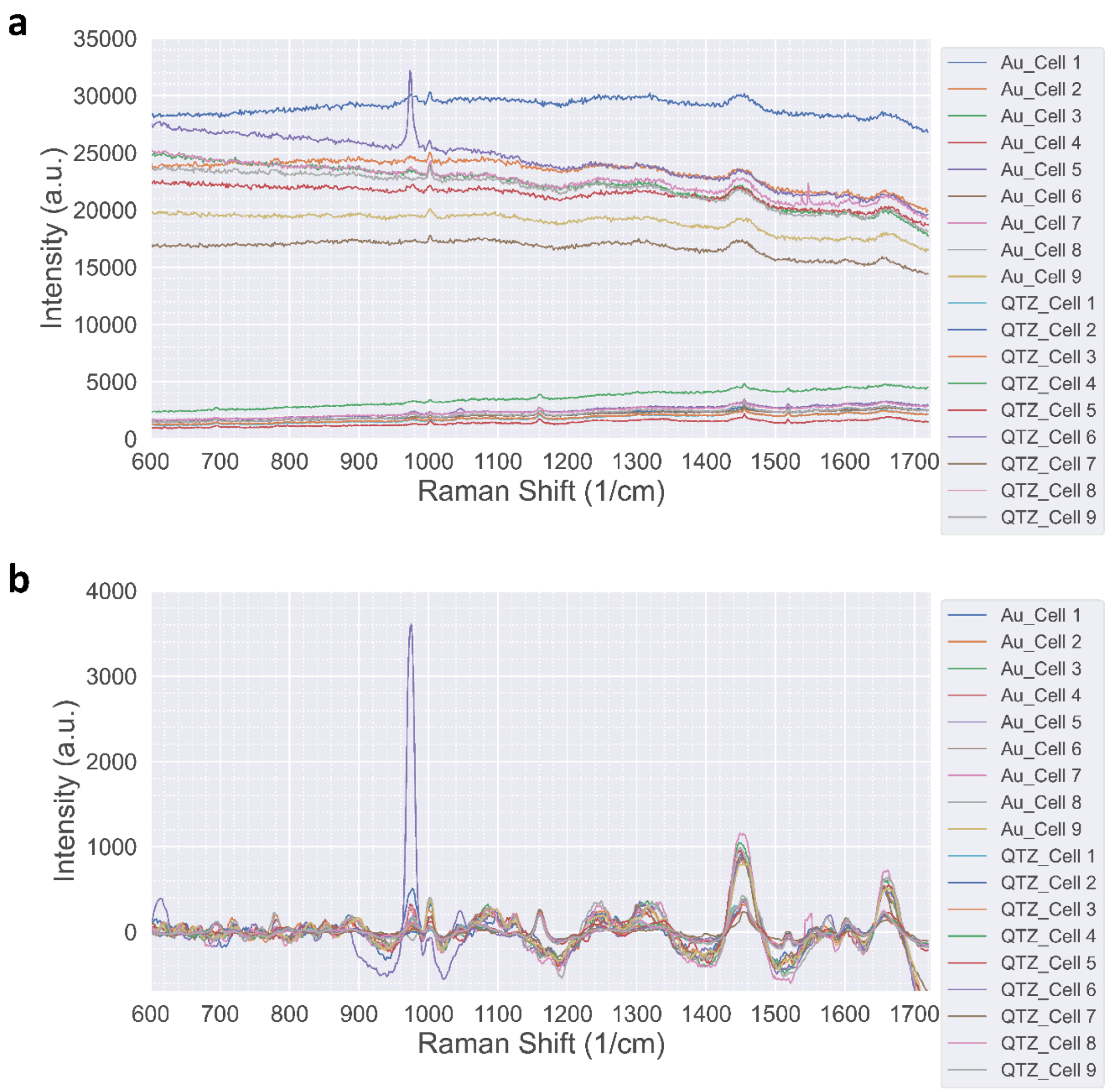


**Figure S5.** Signal amplification of S. cerevisiae Raman spectra using a SERS-active gold substrate. **(a)** Raw Raman spectra of individual S. cerevisiae cells. Spectra acquired from cells on a thin gold (Au) layer exhibit an approximately 10-fold higher signal intensity compared to those from cells on a bare quartz (QTZ) substrate. **(b)** Baseline-corrected spectra from the same cells shown in **(a)**. After background subtraction, the characteristic Raman peaks from cells on the SERS-active gold substrate align with the peaks from cells on bare quartz, validating that the signal is amplified without significant spectral distortion.

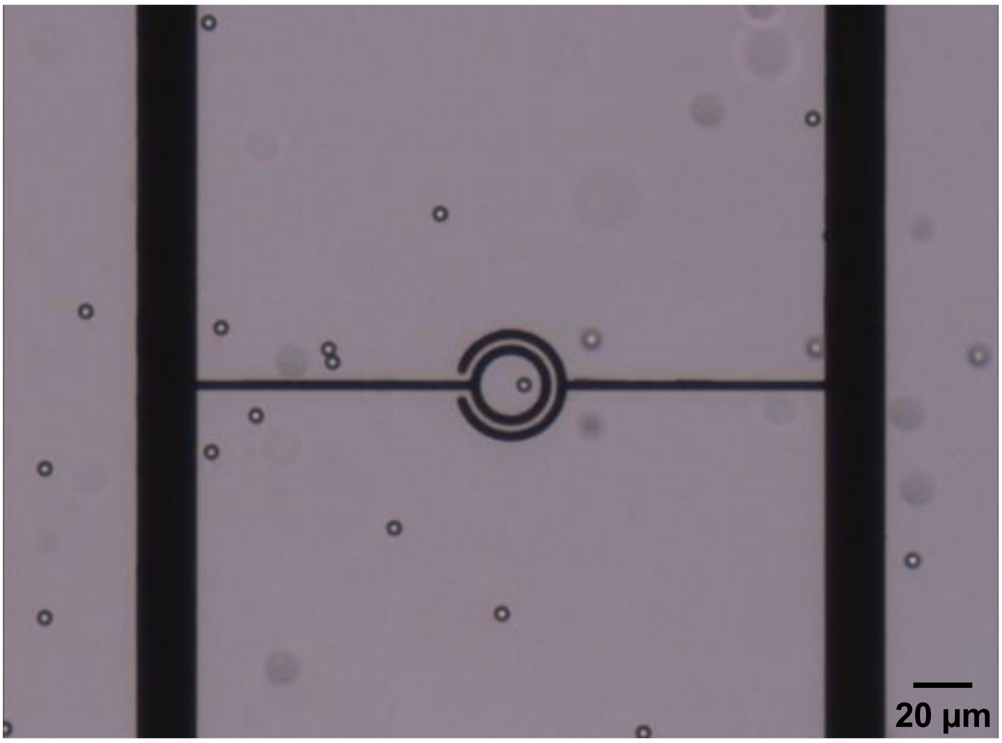


**Movie S1. Trapping of a single 5 µm polystyrene bead.** Here, the bead is trapped under balanced conditions.

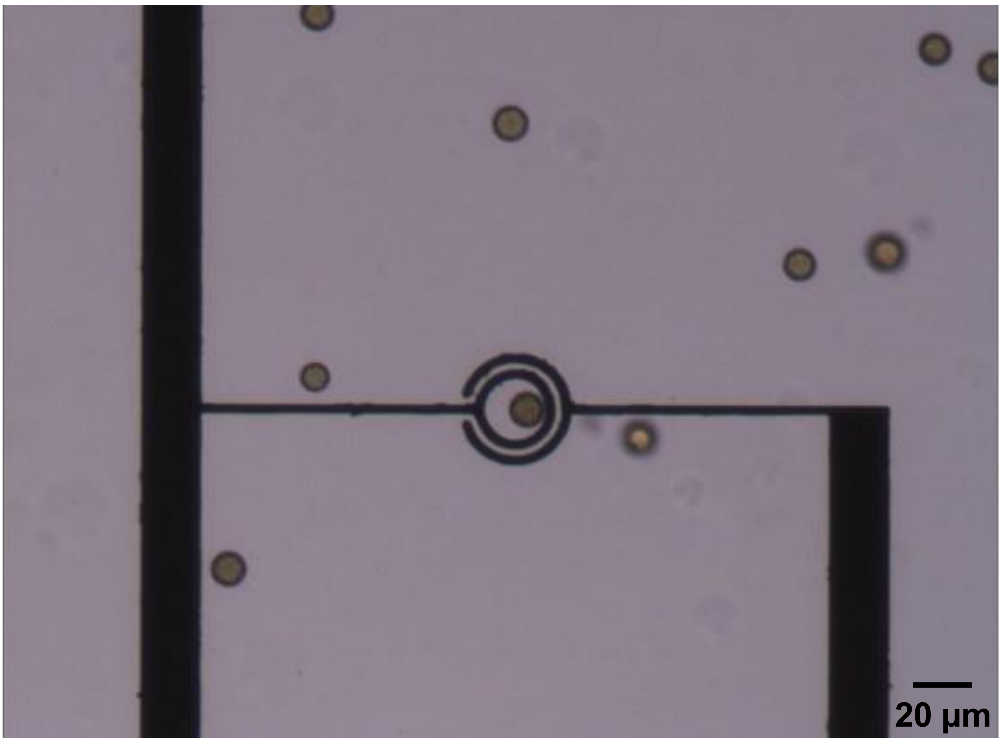


**Movie S2. Trapping of a single 10 µm polystyrene bead.** Here, the bead is trapped under balanced conditions.

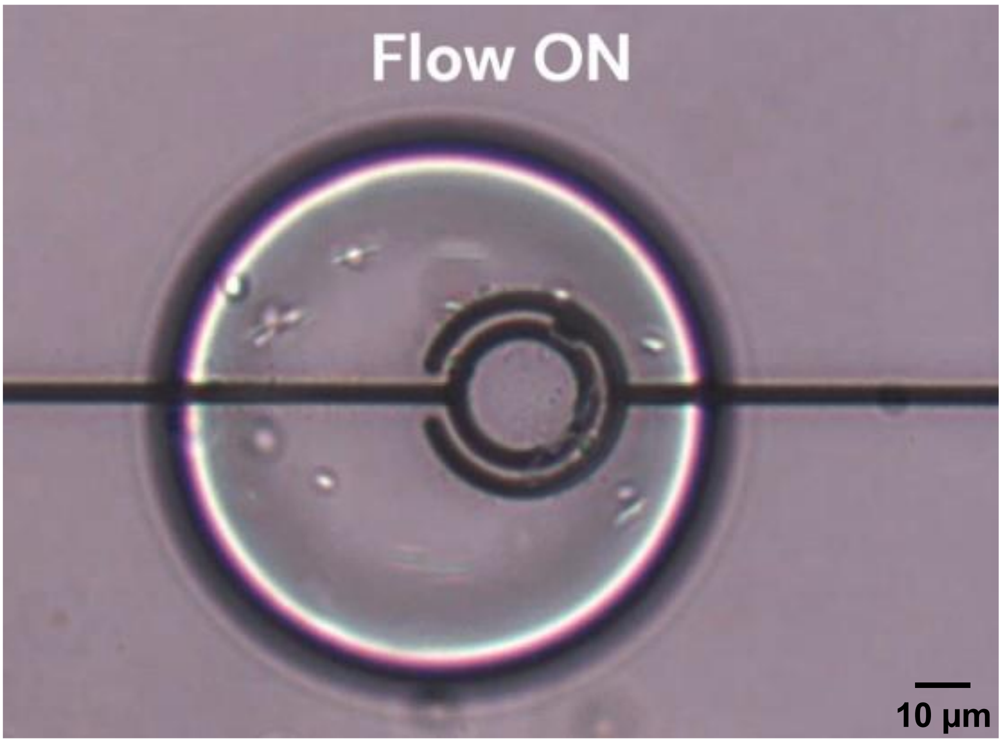


**Movie S3.** Video of a droplet containing *C. albicans* cells under flow. Here, the cells are seen moving freely inside the droplet as it passes the trap without DEP activation.

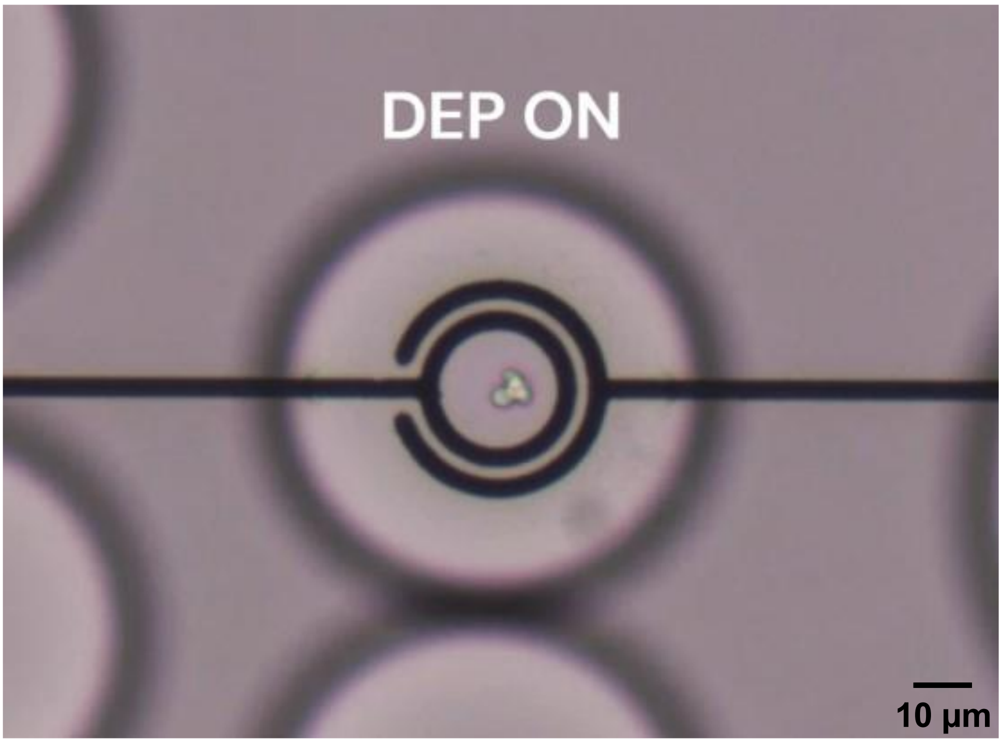


**Movie S4.** Trapping of a droplet containing *C. albicans* cells under no flow. Here, the droplet and the encapsulated cells are immobilized at the nDEP trap upon DEP activation.
